# *O*-GlcNAcylation reduces phase separation and aggregation of the EWS N-terminal low complexity region

**DOI:** 10.1101/2021.05.11.443654

**Authors:** Michael L. Nosella, Maria Tereshchenko, Iva Pritišanac, P. Andrew Chong, Jeffrey A. Toretsky, Hyun O. Lee, Julie D. Forman-Kay

## Abstract

Many membraneless organelles are thought to be biomolecular condensates formed by phase separation of proteins and other biopolymers. Post-translational modifications (PTMs) can impact protein phase separation behavior, although for many PTMs this aspect of their function is unknown. *O*-linked β-D-*N*-acetylglucosaminylation (*O*-GlcNAcylation) is an abundant form of intracellular glycosylation whose roles in regulating biomolecular condensate assembly and dynamics have not been delineated. Using an *in vitro* approach, we found that *O*-GlcNAcylation reduces the phase separation propensity of the EWS *N*-terminal low complexity region (LCR_N_) under different conditions, including in the presence of the arginine-and glycine-rich RNA-binding domains (RBD). *O*-GlcNAcylation enhances fluorescence recovery after photobleaching (FRAP) within EWS LCR_N_ condensates and causes the droplets to exhibit more liquid-like relaxation following fusion. Following extended incubation times, EWS LCR_N_+RBD condensates exhibit diminished FRAP, indicating a loss of fluidity, while condensates containing the *O*-GlcNAcylated LCR_N_ do not. In HeLa cells, EWS is less *O*-GlcNAcylated following *OGT* knockdown and more prone to aggregation based on a filter retardation assay. Relative to the human proteome, *O*-GlcNAcylated proteins are enriched with regions that are predicted to phase separate, suggesting a general role of *O*-GlcNAcylation in regulation of biomolecular condensates.

Insert Table of Contents artwork here

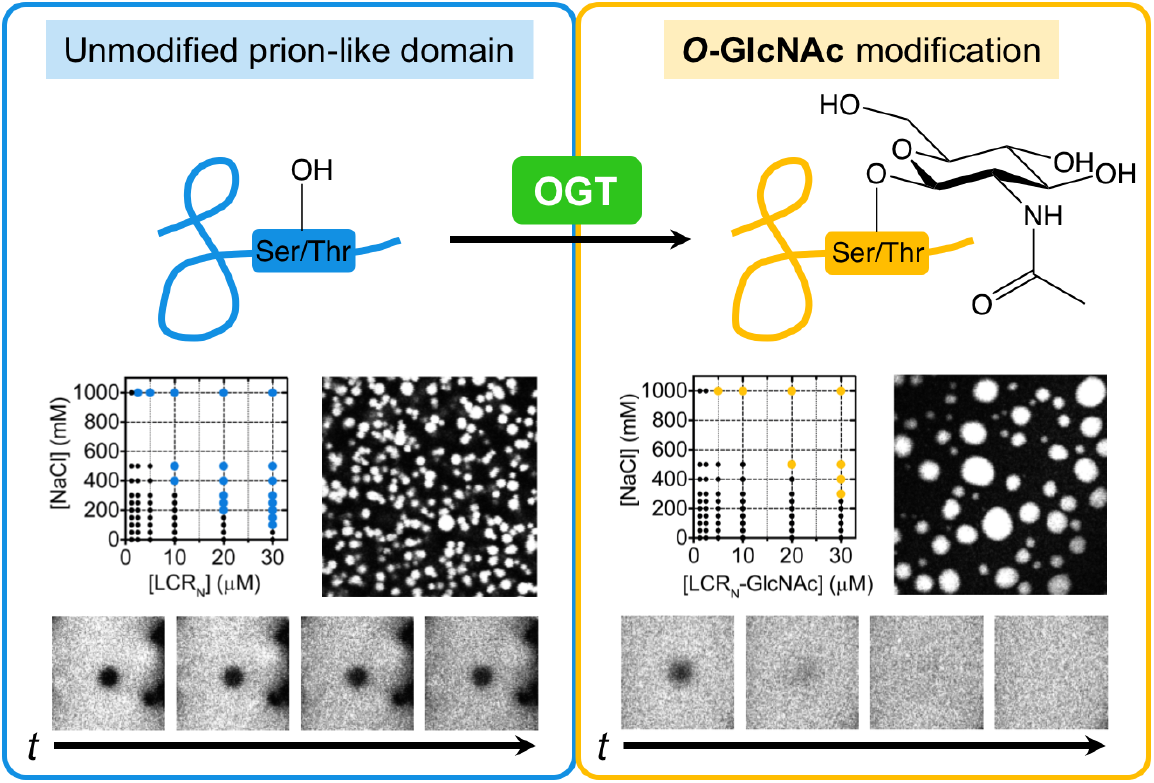

For Table of Contents only.

## INTRODUCTION

The intracellular spaces of living cells are organized into sub-compartments, many of which lack an encapsulating lipid membrane. Several of these ‘membraneless’ compartments are now understood to be condensates formed by phase separation of their constituent biomolecules ^1-3^. Phase-separated condensates can provide unique physical and chemical environments which may contribute to the regulation of diverse biological processes. Determining which mechanisms exist to modulate condensate properties is therefore crucial for understanding their functional (or pathological) roles.

Biological polymers such as proteins and nucleic acids undergo phase separation due to cohesive, multivalent inter-molecular interactions. The numbers and types of interactions derive from physicochemical sequence properties, which, in the case of proteins, are further enrichened by post-translational modifications (PTMs) ^4, 5^. PTMs can impact various aspects of condensates, ranging from composition to larger-scale morphological properties and formation/disassembly kinetics ^6-8^. Consistent with a role for PTMs in modulation of phase separation, the numbers of phosphorylation and methylation sites occurring on proteins statistically correlate with their likelihood to phase separate due to interactions involving pi-contacts ^9, 10^. While PTMs play important and nuanced roles in regulating biological phase separation, for many PTMs this aspect of their function has remained unexplored.

*O*-linked β-D-*N*-acetylglucosaminylation (‘*O*-GlcNAc’ylation) is a form of glycosylation occurring in the nucleus, mitochondria and cytoplasm of higher eukaryotic cells ^11, 12^. Distinct from the often polymeric and structurally complex extracellular glycans, intracellular *O*-GlcNAcylation entails the reversible modification of serine and threonine *O*-hydroxyl groups with a single GlcNAc moiety. In humans and most other metazoa, addition and removal of *O*-GlcNAc are catalyzed by a single pair of enzymes, *O*-GlcNAc transferase (OGT) and *O*-GlcNAc hydrolase (OGA), respectively, which together regulate modification of over a thousand proteins (currently known) with diverse functions ^11, 13-18^. Although *O*-GlcNAcylation is enriched in cellular condensates, such as stress granules ^19^ and DNA damage puncta ^20^, its roles in regulating the formation and physical properties of these and potentially other condensates are poorly understood.

*O*-GlcNAcylation has been found to reduce aggregation and affect phase behavior of certain proteins. For example, *O*-GlcNAcylation of the *Drosophila melanogaster* Polyhomeotic protein enables its incorporation into functional Polycomb repressive complexes by repressing aggregation of its sterile alpha motif domain ^21^. *O*-GlcNAcylation of the nuclear pore complex enhances its passivity and selectivity by tempering interactions between FG-Nups which are otherwise prone to aggregation ^22^. Furthermore, *O*-GlcNAcylation of alpha-synuclein ^23, 24^ and tau ^25^ inhibits their self-association into pathological fibrillar states commonly associated with neurodegenerative disorders.

The connection between phase separation and protein aggregation is highlighted by recent reports of liquid-like condensates preceding (or promoting) aggregate states through liquid-to-solid phase transitions ^26-29^, particularly in the case of RNA-binding proteins (RBPs) such as FUS ^30^ and hnRNPA1 ^31, 32^. Liquid-to-solid phase transitions often rely on regions of low sequence complexity (LCRs), particularly prion-like domains (PLDs) ^30, 33-36^, so-called because of their compositional similarities to bona fide yeast prion proteins ^37^. Perturbations that affect the strengths and/or numbers of protein-protein interactions can affect whether condensates are biased towards more liquid-like or solid-like material states ^30, 38-43^. For example, serine/threonine phosphorylation of the FUS *N*-terminal LCR, which is a PLD, reduces its phase separation propensity as well as its ability to form fibrils by disrupting cohesive cross-beta type interactions between LCR monomers ^40, 44^. Conversely, neurodegenerative disease mutations in FUS and hnRNPA2 can enhance the rate of liquid-to-solid phase transitions by increasing the stabilities of fibrillar structures within initially-liquid condensates ^30, 38^. Fine-tuning of condensates’ physical properties can impact physiological functions as well ^45-48^. Defining how PTMs alter particular condensate material properties is therefore of considerable relevance for understanding condensate function and dysfunction.

The Ewing Sarcoma Breakpoint Region 1 (EWS) protein, a member of the FUS-EWS-TAF15 (FET) protein family that has been frequently examined in the phase separation literature ^33, 41, 49-53^, is known to be *O*-GlcNAcylated *in vivo* ^54^. Given that *O*-GlcNAc reduces protein aggregation and that the EWS N-terminal LCR (LCR_N_) is a PLD with potential for aggregation, we hypothesized that *O*-GlcNAc could modulate the interactions that drive liquid-liquid phase separation of EWS LCRs. Taking an *in vitro* approach, we found that *O*-GlcNAcylation raises the saturation concentration at which the EWS LCR_N_ self-assembles into condensates, including in the presence of the *C*-terminal RNA-binding domains (RBD). *O*-GlcNAcylated EWS LCR_N_ droplets exhibit more liquid-like relaxation and diffusive behavior than un-modified droplets, which behave more like static aggregates. Condensates containing both the EWS LCR_N_ and RBD lose their rapid internal FRAP over time and *O*-GlcNAcylation prevents this. By silencing OGT expression in HeLa cells, we found that lower *O*-GlcNAcylation levels correlated with increased retention of EWS in a filter retardation assay, suggesting that *O*-GlcNAcylation directly affects EWS solubility in cells as it does *in vitro*. From a bioinformatic analysis, we found that a significantly greater fraction of *O*-GlcNAcylated proteins are predicted to phase separate compared to the human proteome as a whole. Based on our results, we propose that *O*-GlcNAcylation could regulate the fluidity and function of LCR-containing biomolecular condensates *in vivo*.

## RESULTS

While all three FET proteins share similar sequence characteristics and cellular localization tendencies, EWS is the sole member shown to be *O*-GlcNAcylated with high stoichiometry *in vivo* ^54^. All three FET proteins contain *N*-and *C*-terminal LCRs (Figure 1A) which undergo phase separation in different contexts, though we did not know if both of these regions in EWS are *O*-GlcNAcylated ^55^. To test the effect of *O*-GlcNAc on EWS phase separation, we purified the LCR_N_ (residues 1-264) and *C*-terminal RBD (residues 265-656) and modified them with recombinant, *Homo sapiens* OGT (nucleocytoplasmic isoform, 110kDa ^56^) using the sugar donor substrate, uridine 5’-diphospho-*N*-acetylglucosamine (UDP-GlcNAc) (Figure 1B). We used electrospray ionization mass spectrometry coupled to liquid chromatography (ESI LC-MS) to assess *O*-GlcNAcylation stoichiometry following the reactions (Figure S1). Following an overnight *O*-GlcNAcylation reaction, LCR_N_ spectra displayed a set of peaks with incremental mass differences of ∼203 Da consistent with discrete *O*-GlcNAc modification states ranging from 3 to 10 moieties per LCR_N_ molecule (Figure 1C, S1A, S1B). The same splitting pattern was not achieved with a catalytically-inactive OGT mutant (K842M), demonstrating a lack of modification in the absence of OGT catalysis (Figure S1C). Spectra of the RBD showed negligible changes, indicating that this region is not *O*-GlcNAcylated *in vitro* (Figure S1D). The distribution of *O*-GlcNAc stoichiometries on the LCR_N_ is consistent with previous reports showing a range of *O*-GlcNAcylation states *in vivo* for EWS ^54, 57^. We conclude that only the *N*-terminal portion of the EWS preceding the RBD is subject to *O*-GlcNAcylation *in vitro*.

**Figure 1.**
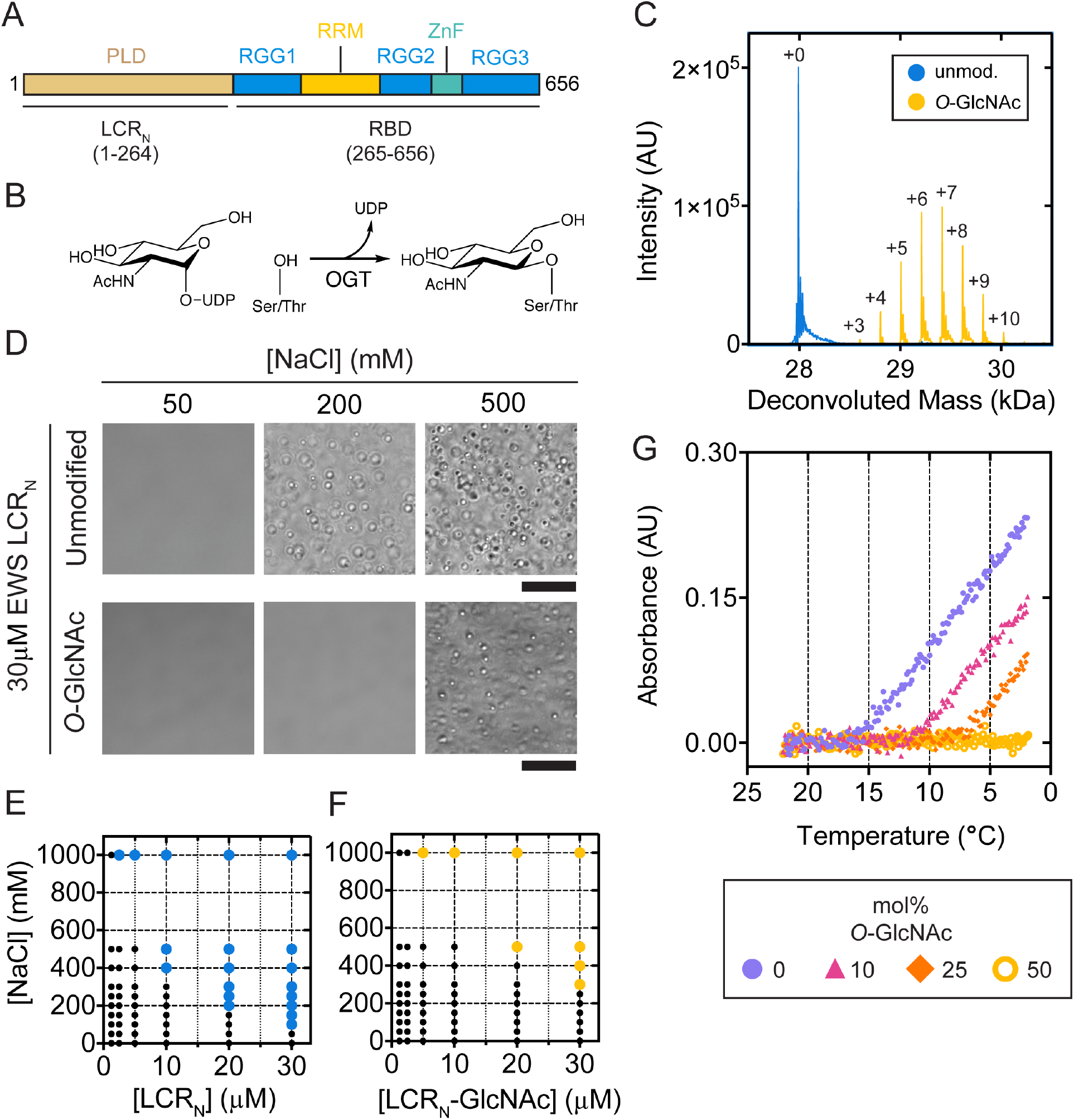
*O*-GlcNAcylation diminishes EWS LCR_N_ phase separation propensity *in vitro*. (A) Schematic of EWS domain organization. PLD, prion-like domain low-complexity disordered region; RGG, arginine-glycine-glycine rich low-complexity disordered region; RRM, RNA-recognition motif; ZnF, zinc finger. (B) Schematic of the OGT-catalyzed *O*-GlcNAcylation of serine and threonine sidechains. (C) Deconvoluted mass spectra corresponding to unmodified (blue) and *O*-GlcNAcylated (yellow) LCR_N_. Labels indicate number of *O*-GlcNAc modifications. (D) Differential interference contrast (DIC) micrographs of 30µM unmodified (top) or *O*-GlcNAcylated (bottom) EWS LCR_N_ in 25mM Tris pH 7.5 at lab temperature with denoted NaCl concentrations. Scale: 20µm. (E,F) Salt and concentration conditions under which (E) unmodified or (F) *O*-GlcNAcylated EWS LCR_N_ formed droplets. Large, colored dots indicate conditions for which droplets were observed; small black dots indicate an absence of droplets. (G) Temperature-dependent turbidity (λ = 500nm) of LCR_N_ samples containing variable proportions of *O*-GlcNAcylated LCR_N_. Total LCR_N_ concentration is consistently 30µM; legend indicates the proportion of *O*-GlcNAcylated LCR_N_ present in the samples.

Like the *N*-terminal LCR of FUS, the EWS LCR_N_ self-associates into hydrogels and induces sedimentation of the chimeric fusion protein EWS-FLI1 *in vitro* ^41, 53^, leading us to hypothesize that the EWS LCR_N_ could also form liquid-like droplets on its own. Addition of the EWS LCR_N_ (30µM) into buffer solutions at pH 7.5 with 500mM NaCl immediately resulted in the formation of round, micron-sized foci (hereafter termed ‘droplets’) which were absent at lower salt concentrations (Figure 1D). Droplet formation correlated with increasing NaCl and LCR_N_ concentrations in a similar manner as the FUS LCR_N_ ^58^ (Figure 1E). To test the influence of *O*-GlcNAc on droplet formation, we purified milligram quantities of the *O*-GlcNAcylated EWS LCR_N_ from a scaled-up OGT reaction (see SI Methods). The *O*-GlcNAcylated LCR_N_ formed droplets with similar morphologies as the unmodified protein, albeit at higher NaCl and protein concentrations (Figure 1E,F), implying that *O*-GlcNAc lowers the propensity of the EWS LCR_N_ to phase separate under these conditions. Like the FUS LCR, we also expected EWS LCR_N_ phase separation to occur at decreasing temperatures in accordance with upper critical solution temperature-type behavior ^58^. We found that unmodified LCR_N_ samples at pH 7.5 in the presence of 100mM NaCl (i.e., below saturating conditions at room temperature) became turbid as the temperature was decreased below ∼16°C, indicating droplet formation below this temperature (Figure 1G circle). Increasing the fraction of *O*-GlcNAcylated LCR_N_ in the samples led to an incremental lowering of the saturation temperature (Figure 1G triangle, diamond). At a 1:1 ratio of *O*-GlcNAcylated:unmodified LCR_N_, turbidity was absent at all points of the temperature series under these conditions (Figure 1G ring). Thus, *O*-GlcNAcylation reduces the propensity of EWS LCR_N_ to phase separate in response to increasing salt concentrations and decreasing temperature.

To determine if phase behavior correlates with the number of *O*-GlcNAc modifications, we prepared small-scale *O*-GlcNAcylation reactions containing the EWS LCR_N_ and varying concentrations of OGT. Following a 30-minute reaction period, the NaCl concentration was adjusted to 200mM to promote droplet formation, which we measured by turbidity. EWS LCR_N_ samples displayed an approximately six-fold increase in turbidity following the addition of NaCl (Figure 2A). Inclusion of both OGT and UDP-GlcNAc resulted in no turbidity change at 30 minutes relative to samples kept at low (50mM) NaCl (Figure 2A). The relative turbidity scaled inversely with the number of *O*-GlcNAc modifications (Figure 2B,C). Substituting wild-type OGT with the catalytically-deficient mutant (K842M) did not produce any changes in turbidity with respect to enzyme concentration (Figure 2B), meaning that the reduction in turbidity seen with wild-type was not merely caused by enzyme binding-induced solubilization of the LCR_N_. Even the lowest tested OGT concentrations (0.05, 0.1µM) resulted in slight reductions in turbidity, despite only a minor fraction of LCR_N_ molecules being *O*-GlcNAcylated under these conditions (Figure 2C). These data suggest that *O*-GlcNAcylation lessens salt-induced phase separation of the EWS LCR_N_ even at sub-stoichiometric modification levels. We then asked whether the LCR_N_ can be *O*-GlcNAcylated in the condensed phase after phase separation, and if so, will *O*-GlcNAcylation cause the droplets to disassemble. We performed turbidity assays in which LCR_N_ phase separation was induced with 200mM NaCl prior to the onset of *O*-GlcNAcylation. Following a 30-minute reaction period, LCR_N_ samples containing OGT and UDP-GlcNAc yielded similar modification patterns to those for reactions per formed in the dilute phase (Figure S1E). All samples, including those lacking either OGT or UDP-GlcNAc, displayed time-dependent decreases in turbidity due to the droplets sinking. However, the samples containing both OGT and UDP-GlcNAc underwent a significantly greater decrease over the 30-minute time course, after which few droplets could be observed by microscopy (Figure 2D,E). Samples lacking either OGT or UDP-GlcNAc contained droplets before and after the reaction period (Figure 2D,E). Interestingly, we never witnessed total clearance of the droplets, perhaps because levels of *O*-GlcNAcylation were generally lower than when the LCR_N_ was modified in the dilute state (Figure S1E). This could have resulted from the non-mutually exclusive possibilities of OGT having lower catalytic turnover rates in the presence of 200mM NaCl or in the droplets, or from its lower partitioning into the condensed phase. These data demonstrate that *O*-GlcNAcylation can effectively reverse EWS LCR_N_ droplet formation in vitro.

**Figure 2.**
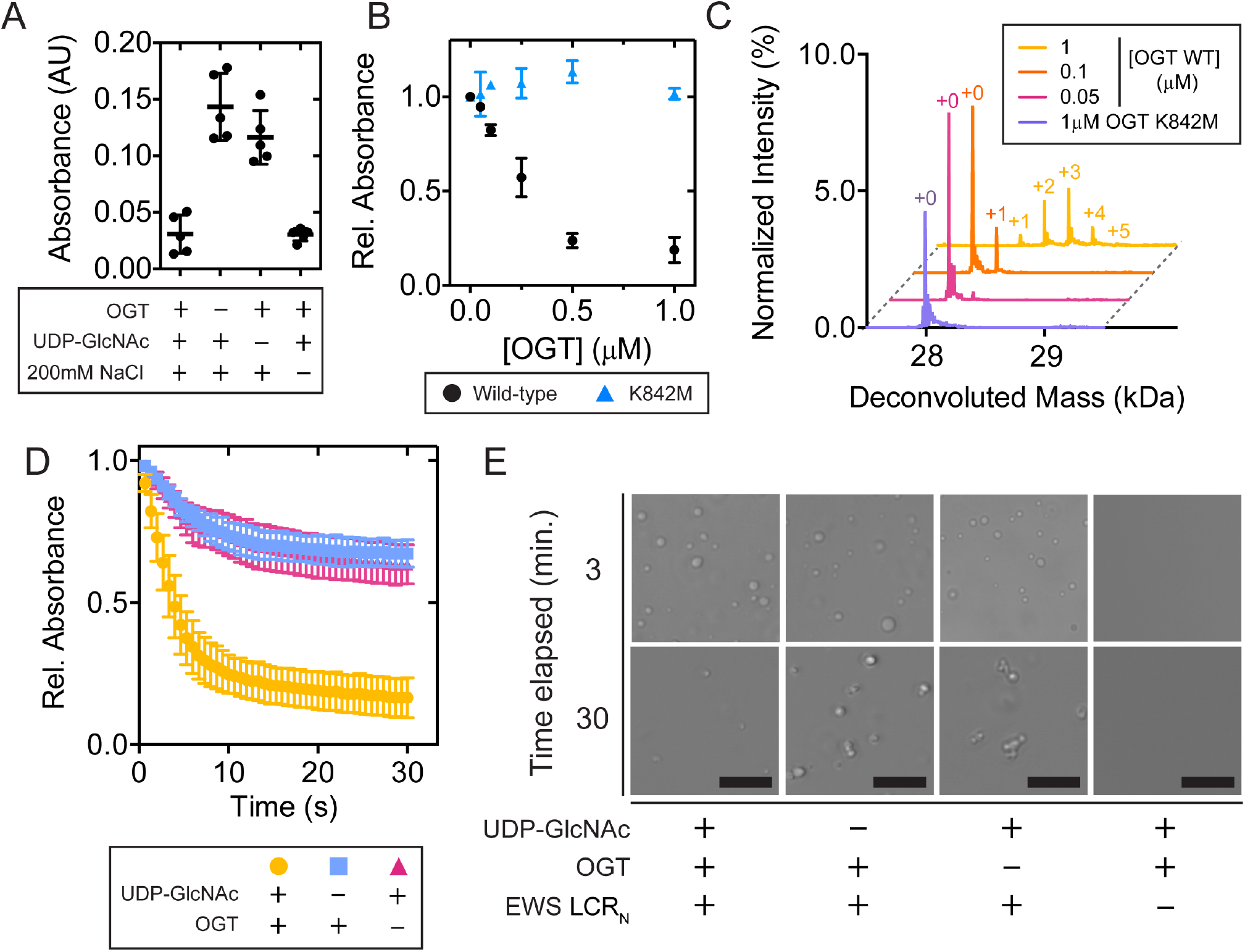
EWS LCR_N_ phase separation propensity is reduced as *O*-GlcNAc modifications increase. (A) Turbidity (λ = 500nm) of LCR_N_ (20µM) *O*-GlcNAcylation reaction samples following addition of NaCl. Central bold bars represent mean, with error bars showing standard deviation, n = 5. (B) Relative turbidity (λ = 500nm) of LCR_N_ (20µM) *O*-GlcNAcylation reactions as a function of varying OGT wild-type (black) or K842M (blue) concentration. Droplet formation was induced by addition of NaCl following a 30-minute reaction period. Error bars represent standard deviation, n = 3. (C) LC-ESI MS analysis of samples in (B). (D) Relative turbidity (λ = 500nm) of LCR_N_ *O*-GlcNAcylation reactions containing phase-separated EWS LCR_N_ (20µM) as a function of reaction time. Errors bars represent standard deviation, n = 5. (E) DIC micrographs corresponding to samples in (D) at 3 or 30 minutes post-addition of UDP-GlcNAc. Scale: 20µm. Unless stated otherwise, OGT and UDP-GlcNAc concentrations in each reaction were 1µM and 1mM, respectively.

We then examined how *O*-GlcNAcylation affects the morphologies of EWS LCR_N_ droplets over time. Within minutes after formation, the unmodified LCR_N_ droplets cluster into bleb-like structures with irregular boundaries, whereas the *O*-GlcNAcylated droplets merge into larger, circular droplets with smooth boundaries (Figure 3A). To test if the *O*-GlcNAc-dependent morphological differences were caused by changes in the droplets’ material properties, we measured the impact of *O*-GlcNAc on droplet fusion, which is a diagnostic characteristic of liquid-like behavior in protein condensates. We prepared samples of the unmodified and *O*-GlcNAcylated LCR_N_ droplets spiked with 100nM (∼0.3mol%) sulfo-cyanine-3 (sulfo-Cy3)-labelled protein for visualization by scanning confocal fluorescence microscopy (Figure 3A). After colliding, *O*-GlcNAcylated droplets spontaneously merged and relaxed into circular shapes when viewed along the xy-plane, tending to a unitary aspect ratio within seconds to minutes (Figure 3B,C). The unmodified droplets adhered upon collision but did not deform, instead retaining their pre-fusion morphologies. For the *O*-GlcNAcylated droplets, the exponential decay in aspect ratio was linearly proportional to the length scale of the fusion event, consistent with previously reported behavior of liquid-like protein condensates in vitro (Figure 3D) ^59^. *O*-GlcNAcylated EWS LCR_N_ droplets fuse more slowly than other in vitro protein condensates such as those formed by NPM1 46, LAF-1 ^60^ or PR_60_-repeat peptides with RNA ^61^, perhaps indicating higher viscosity or surface tension, though the comparison may be skewed by differences in sample conditions.

**Figure 3.**
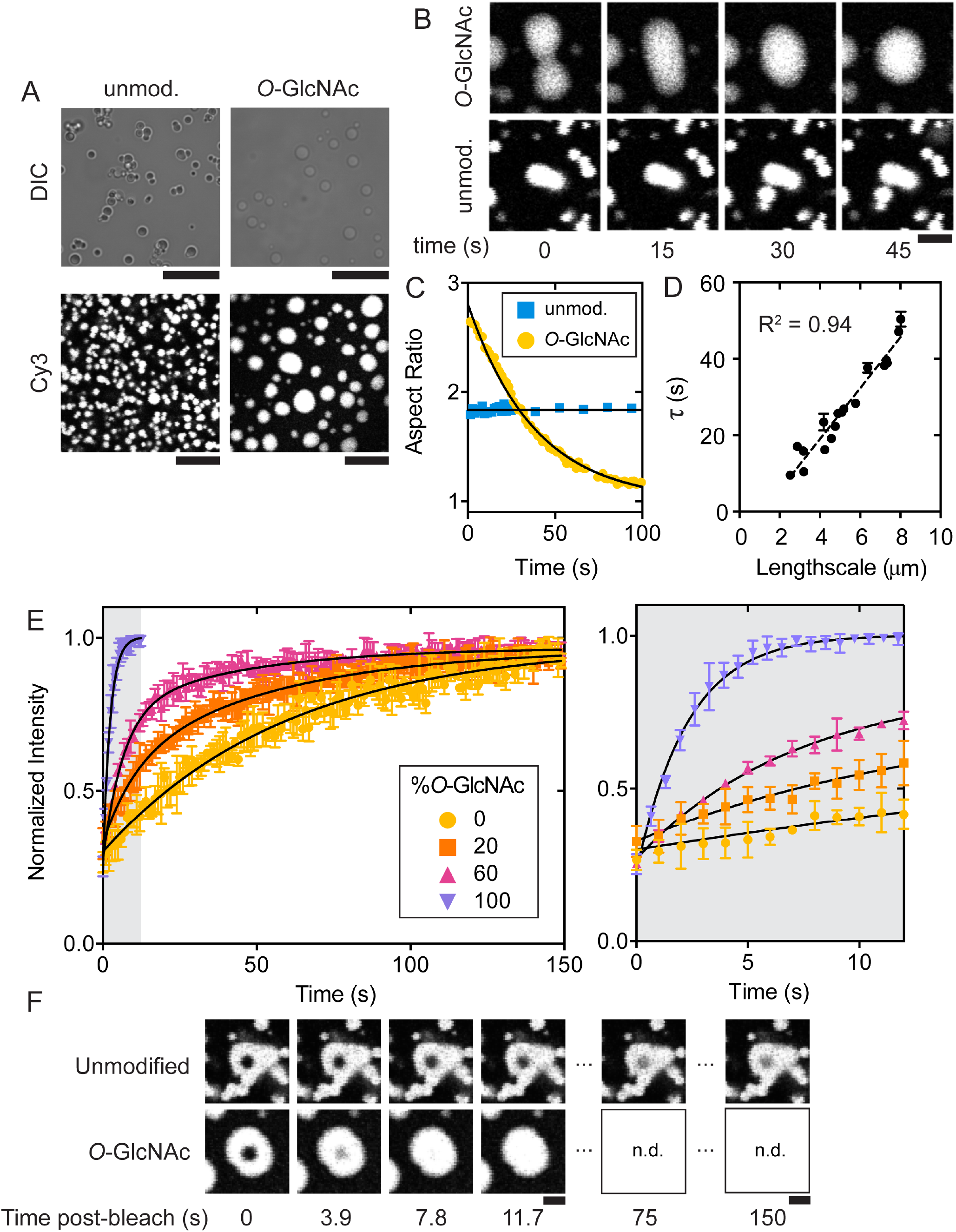
*O*-GlcNAcylation enhances fluidity of EWS LCR_N_ droplets. (A) DIC and fluorescence micrographs of unmodified (left) or *O*-GlcNAcylated (right) LCR_N_ droplets. Droplets spiked with 100nM sulfo-Cy3-labelled LCR_N_ (with or without *O*-GlcNAcylation) were imaged at 30 min. (middle) or 90 min. (bottom) post-formation. Both samples contain 30µM total protein concentration, in 25mM Tris, 500mM NaCl, pH 7.5 at lab temperature. Scale: 20µm. (B) Time lapse images of unmodified (bottom) and *O*-GlcNAcylated (top) sulfo-Cy3-spiked LCR_N_ droplets fusing (same conditions as in (A)). Scale: 5µm. (C) Plot of droplet aspect ratio versus time for representative fusion events displayed in (B). (D) Time constants (τ) of exponentially decaying *O*-GlcNAcylated droplet aspect ratio versus length scale (distance between fusing droplets). The slope of the linear fit estimates the inverse capillary velocity (ratio of effective droplet viscosity to surface tension): ∼6.6µm s^-1^. Error bars represent the SEM of fitted τ values. (E) FRAP traces from droplets containing variable ratios of *O*-GlcNAcylated:unmodified EWS LCR_N_, spiked with 100nM sulfo-Cy3-labelled LCR_N_ (with or without *O*-GlcNAcylation). *Right*: Zoom-in of grey shaded area in left panel. N = 4; error bars represent standard deviation. (F) Fluorescence micrographs of representative FRAP time points in (E). n.d.: no data. Scale: 5µm.

To determine whether the diffusion of LCR_N_ molecules within droplets is affected by *O*-GlcNAcylation, we performed fluorescence recovery after photobleaching (FRAP) measurements on the same sulfo-Cy3-spiked droplet samples. Within the unmodified droplets, fluorescence intensity returned to ∼90% of pre-bleach levels within a 2.5µm-diameter photobleached region in under 3 minutes (Figure 3E,F), indicating that the majority of molecules remain mobile despite the droplets’ inability to fuse. Applying the same photobleaching parameters onto the *O*-GlcNAcylated LCR_N_ droplets resulted in similar levels of fluorescence recovery on a significantly reduced time scale (Figure 3E,F). Fitting a monophasic exponential model to these data gave fluorescence recovery half-times of 41.6 ± 1.7s and 1.59 ± 0.07s for the unmodified and *O*-GlcNAcylated droplets, respectively. In droplets containing mixed ratios of O-GlcNAcylated and unmodified LCR_N_, the recovery half-times decreased proportionally with the amount of *O*-GlcNAcylated LCR_N_ present (Figure 3E). We cannot confidently estimate diffusion coefficients from these data due to the dissimilar sizes of the differentially modified droplets as well as the potential flaws in using a single-exponential model to describe the underlying diffusion mechanisms ^62^. However, the difference in half-times is striking, clearly demonstrating that *O*-GlcNAcylation enhances diffusion of EWS LCR_N_ molecules within condensates. Based on the disparate fusion behaviors and accelerated FRAP, we suggest that *O*-GlcNAcylation enhances the liquidity of EWS LCR_N_ droplets, which are otherwise highly viscous or perhaps gel-like in the absence of this modification.

FET protein phase separation is not solely driven by LCR_N_ self-interactions; rather, the tyrosine-rich LCR_N_ and arginine-rich RBDs synergistically promote phase separation ^9, 50, 63^. Thus, we sought to determine if *O*-GlcNAcylation would impact co-phase separation of these two regions of EWS (Figure 1A). We titrated the EWS LCR_N_ with or without *O*-GlcNAcylation into samples of the EWS RBD under low-salt buffer conditions in which neither would phase separate on its own. Turbidity increased proportionally with the concentration of unmodified LCR_N_ and coincided with the presence of droplets (Figure 4A,B). We saw a similar increase as we titrated the *O*-GlcNAcylated LCR_N_, however, the onset of turbidity occurred at a higher LCR_N_ concentration, indicating lower phase separation propensity (Figure 4A,B). This effect was more pronounced when the level of *O*-GlcNAcylation was increased, supporting a direct link between *O*-GlcNAc levels and RBD-dependent droplet formation. As a point of comparison, we also tested the effect of asymmetric dimethylation of the RBD arginines (another PTM shown to mitigate phase separation of FUS ^52, 63^) under the same conditions, finding that the increase in saturation concentration was approximately the same as that caused by LCR_N_ *O*-GlcNAcylation (Figure 4A).

**Figure 4.**
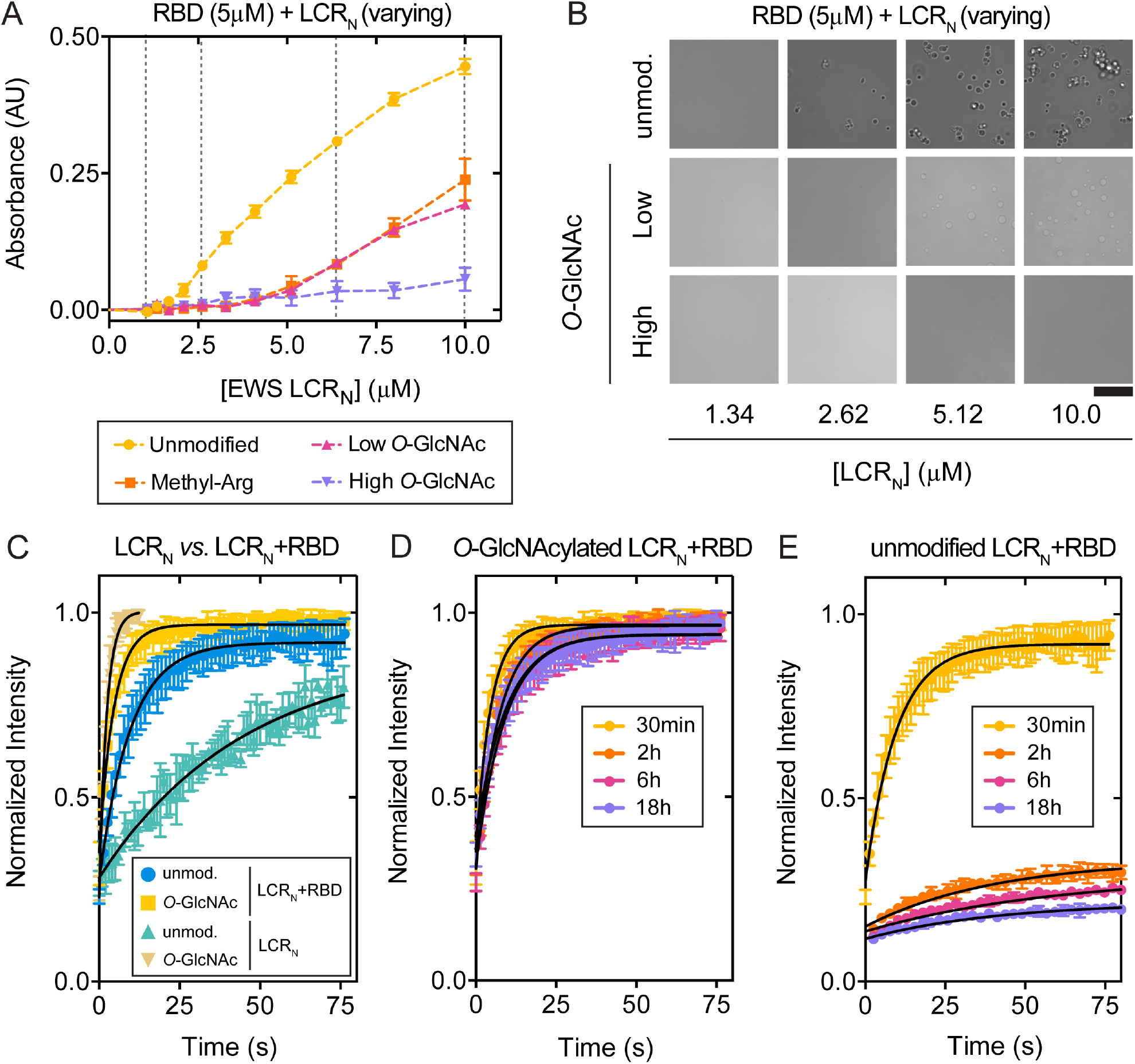
EWS LCR_N_ *O*-GlcNAcylation reduces RBD-induced phase separation and prevents time-dependent loss of droplet dynamics. (A) Turbidity (λ = 500nm) of EWS RBD samples (5µM; with or without arginine methylation) titrated with LCR_N_ (with or without *O*-GlcNAcylation) in 25mM Tris, 100mM NaCl, 0.5mM tetrasodium EDTA, 2mM dithiothreitol, pH 7.5 at 25°C. Turbidity was measured immediately after mixing of the two EWS fragments. Error bars represent standard deviation, n = 3. (B) DIC micrographs of samples in (A) indicated by hatched vertical lines. Scale: 20µm. (C) FRAP traces for EWS LCR_N_ and LCR_N_+RBD droplets with and without LCR_N_ *O*-GlcNAcylation. LCR_N_ droplet samples are the same as in (3E). LCR_N_+RBD droplet samples contain 10µM LCR_N_, 100nM sulfo-Cy3-labelled LCR_N_, 10µM RBD, 25mM Tris, 100mM NaCl, 0.5mM tetrasodium EDTA, 2mM dithiothreitol, pH 7.5 at lab temperature. (D) FRAP traces of *O*-GlcNAcylated and (E) unmodified LCR_N_+RBD droplets measured at different time points following phase separation under the same conditions as (C). Error bars represent standard deviation, n = 3. Black lines are single exponential fits.

We then tested the effect of LCR_N_ *O*-GlcNAcylation on the diffusive characteristics of droplets containing both the LCR_N_ and RBD (LCR_N_+RBD droplets), spiked with sulfo-Cy3-LCR_N_. Interestingly, the unmodified LCR_N_+RBD droplets exhibited more rapid fluorescence recovery than the unmodified LCR_N_ droplets without addition of the RBD 4C).

While the FRAP kinetics of the unmodified and *O*-GlcNAcylated LCR_N_+RBD droplets were similar within 30minutes following phase separation, they began to diverge at later time points (Figure 4D,E). *O*-GlcNAcylated LCR_N_+RBD droplets exhibited near-complete fluorescence recovery irrespective of the incubation time (Figure 4D,S2), whereas the unmodified droplets already showed decreasing fluorescence recovery at 2 hours post-formation (Figure 4E,S2). The mobile fraction in the unmodified LCR_N_+RBD droplets decreased even further at later timepoints, reaching only ∼10% after 18 hours. These results suggest that molecules within the droplets containing the unmodified LCR_N_ lose the ability to diffuse in a time-dependent fashion indicating a progression to a less fluid-like material state. *O*-GlcNAcylation prevents this loss of mobility, instead maintaining the liquid-like droplet dynamics that are seen shortly after phase separation.

Based on our observations *in vitro*, we investigated the effect of *O*-GlcNAcylation on EWS aggregation propensity in cells. We used a filter retardation assay (FRA) to monitor EWS aggregation in HeLa cells following changes in *O*-GlcNAcylation levels. In this assay, large (>0.2µm) sodium dodecyl sulfate (SDS)-insoluble protein aggregates are captured by vacuum filtration on a cellulose-acetate membrane, while smaller protein assemblies and monomers pass through. The captured aggregates are subsequently detected by immunoblotting. To modulate *O*-GlcNAcylation levels, we downregulated OGT expression using small interfering RNA (siOGT), which caused a global reduction in *O*-GlcNAcylation levels (Figure S3A) including reduction in *O*-GlcNAcylation of EWS as measured by chemoenzymatic labeling (Figure 5A). We found that siOGT treatment led to increased EWS aggregation signal in the FRA versus untreated control samples (Figure 5B,C). In contrast, TAF15, another FET family member that is known to phase separate and/or aggregate ^36, 50, 64^ but is not *O*-GlcNAcylated 54 (Figure S3B), did not experience greater aggregation following siOGT treatment (Figure 5B,C). This suggests that the increased EWS aggregation is a direct consequence of its *O*-GlcNAcylation status rather than a global, protein-nonspecific aggregation caused by decreasing *O*-GlcNAcylation levels. OGT knockdown did not significantly affect the expression levels of EWS or TAF15 (Figure S3C), which could contribute to either protein’s aggregation. Taken together, these results show that the aggregation propensity of EWS is directly correlated with its *O*-GlcNAcylation status in cells.

**Figure 5.**
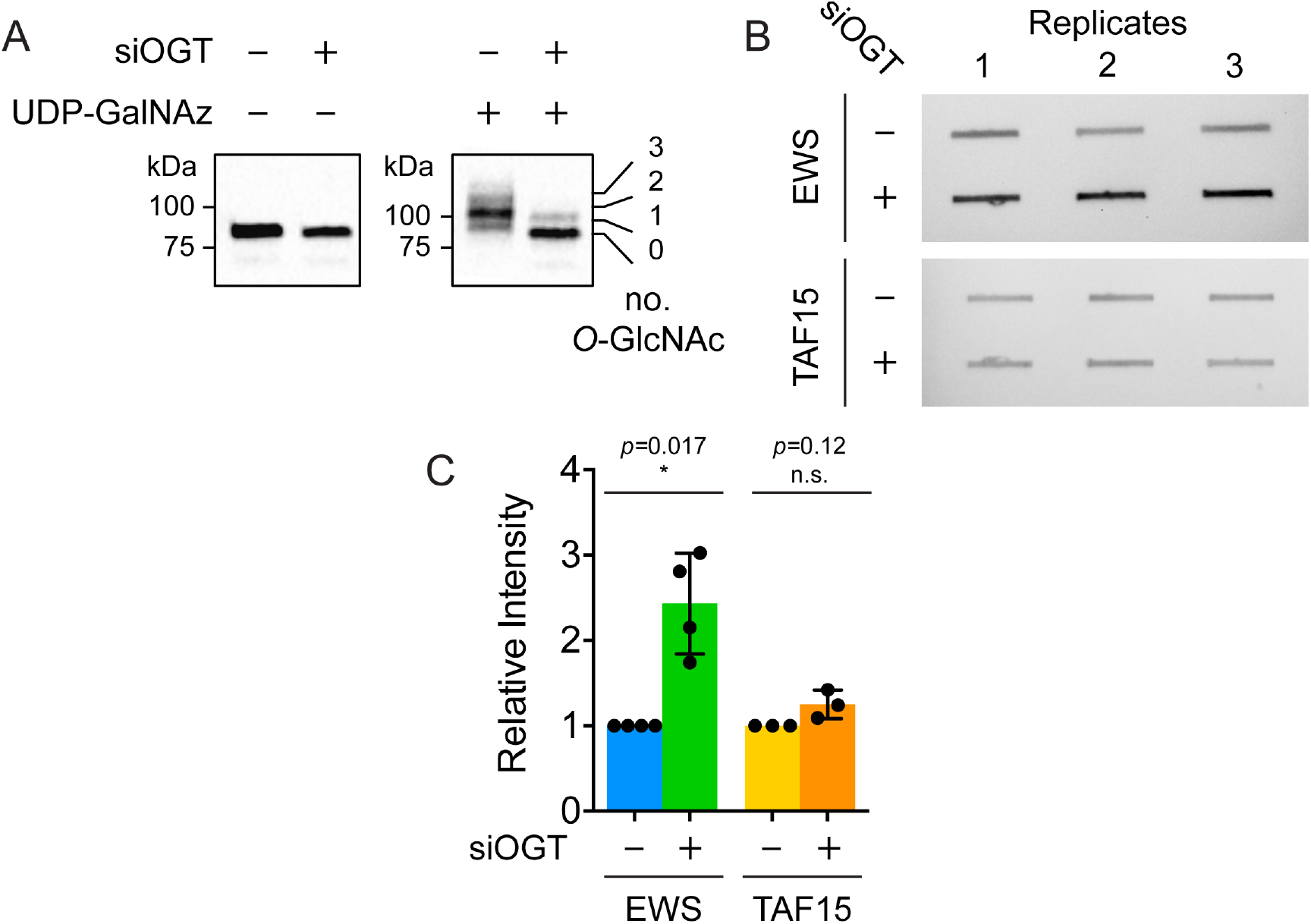
*O*-GlcNAcylation decreases EWS aggregation in HeLa cell lysates. (A) Western blot (anti-EWS) depicting chemoenzymatic (ClickIT) labelling of *O*-GlcNAc moieties on EWS isolated from HeLa cells, treated with or without siOGT (48-hour incubation with 20nM small interfering RNA for OGT). GlcNAc moieties are enzymatically modified with uridine-diphospho-*N*-azidoacetylgalactosamine (UDP-GalNAz) and then alkynyl-PEG (average molecular weight: 5kDa) via Huisgen cycloaddition (‘Click’ chemistry) to produce an apparent molecular weight shift of ∼5kDa per GlcNAc moiety via SDS-PAGE. (B) Representative images of FRA membranes following Western blot detection of EWS (top) or TAF15 (bottom), with and without siOGT treatment. (C) Normalized EWS and TAF15 Western blot intensities from three FRA replicates performed with OGT-depleted HeLa cell lysates. Error bars represent standard deviation. For EWS: *p* = 0.017, Student’s *t*-test; n = 4. For TAF15: *p* = 0.12, Student’s *t*-test; n = 3.

Supposing that *O*-GlcNAcylation might be used by the cell to inhibit aggregation of some phase-separating proteins, we then asked if *O*-GlcNAcylation is generally correlated with higher phase separation propensity. We performed a bioinformatic analysis to calculate the percentage of *O*-GlcNAcylated human proteins predicted to phase separate by three metrics, PScore ^9^, catGRANULE ^65^ and PLAAC ^66, 67^, compared to that for all post-translationally modified proteins included in the PhosphoSitePlus PTM database and the entire human proteome. *O*-GlcNAcylated proteins show a significant enrichment for increased phase separation propensity for each of the three metrics as defined by the fraction scoring above each predictor’s preestablished threshold (Figure S4A). To account for differences in the number of proteins above each predictor’s cut-off, we also compared the fractions that scored in the top 4th percentile, finding that the enrichment in *O*-GlcNAcylated proteins remains consistent (Figure 6). The significant enrichment also persists across the metrics when querying for proteins with long (>100 residue) IDRs (Figure S4B), indicating that the potentially greater proportion of *O*-GlcNAc site annotations being present in long IDRs and the greater proportion of phase-separating proteins having long IDRs does not affect this finding. These results suggest that phase-separating proteins are more likely to be subject to *O*-GlcNAc-mediated regulation than the proteome in general. PLAAC, which predicts protein regions with prion-like sequence compositional bias ^66^, showed the greatest statistical enrichment of the three metrics, implying that *O*-GlcNAcylation may be especially relevant for PLD-containing, phase-separating proteins that are prone to aggregation.

**Figure 6.**
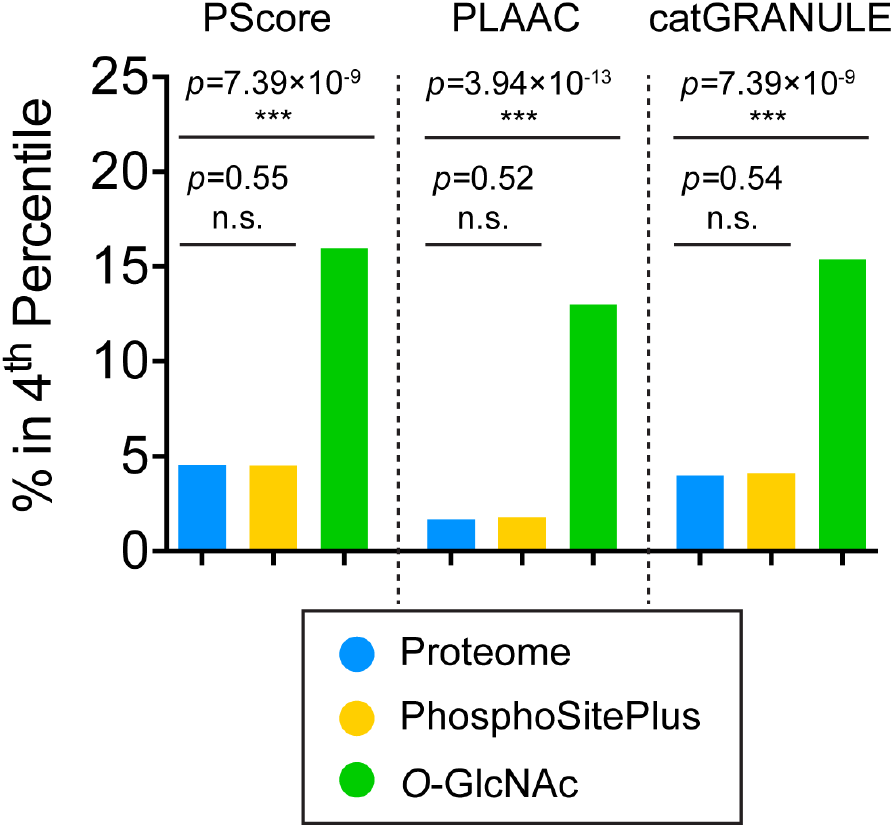
O-GlcNAcylated proteins are enriched for predicted phase separation propensity. Comparison of proteins scoring in the top 4^th^ percentile of the proteome by three different metrics of phase separation propensity (PScore, PLAAC and catGRANULE). Protein sequences for the human proteome were retrieved from the Uniprot database (n = 21047, blue). The human proteins with documented post-translational modifications (n = 19990, yellow) and those modified through *O*-GlcNAcylation (n = 171, green) were retrieved from the PhosphoSitePlus database. The *O-*GlcNAcylated proteins show a significant enrichment for increased phase separation propensity relative to the proteome and post-translationally modified human proteins for each of the three metrics. Indicated *p*-values were derived from Fisher’s exact test.

## DISCUSSION

We examined the effects of *O*-GlcNAc, a widespread PTM with diverse functional consequences ^11, 15, 16^, on the phase separation and material properties of EWS condensates. *O*-GlcNAcylated LCR_N_ requires higher protein and salt concentrations and lower temperatures to form droplets, evidence for a decreased phase separation propensity. *O*-GlcNAcylated EWS droplets are more fluid than unmodified droplets based on their increased relaxation to spherical shapes following fusion and their more rapid FRAP. Inclusion of the EWS RBD in trans promotes fluidity at short times, although the internal mobility of LCR_N_+RBD droplets diminishes as the unmodified droplets age, perhaps due to a time-dependent increase in PLD-induced cross-beta structure 33. This transition does not occur when the LCR_N_ is *O*-GlcNAcylated, suggesting that *O*-GlcNAc preserves fluid-like condensate dynamics over time.

The effects of *O*-GlcNAcylation on the assembly and dynamics of higher-order protein complexes has already been described in several systems, including FG-Nup hydrogels ^22^, Drosophila Polycomb complexes ^21^, intermediate filaments ^68^, and pathological fibrillar aggregates formed by tau and alpha-synuclein ^23-25^. In each of these cases, *O*-GlcNAcylation tends to promote protein solubility by sterically disrupting intermolecular interactions between monomers. Our results suggest that the same effect may apply to EWS LCR_N_ condensates, whose phase separation and liquid-like dynamics are sensitive to the strengths and numbers of *O*-GlcNAc-modulated intermolecular interactions. We demonstrate that *O*-GlcNAc modulates EWS condensate dynamics even at sub-stoichiometric levels, meaning that not all molecules within the condensate must be modified in order for bulk condensate properties to be affected. These results mirror previous findings that sub-stoichiometric *O*-GlcNAcylation slows formation of alpha-synuclein fibrils ^23^. Thus, *O*-GlcNAcylation may influence condensate properties even when the modified species are not the most abundant components of the system.

Interactions between tyrosine and arginine residues are significant drivers of phase separation of PLD-containing RNA-binding proteins such as EWS ^50^. *O*-GlcNAc reduces phase separation of the tyrosine-rich EWS LCRN together with the arginine-rich RBD, suggesting that it can affect their interactions even though neither of these key residue types ^50^ are directly modified. The sugar may sterically block cohesive interactions important for phase separation and aggregation, including those underlying cross-beta structures ^40^. In addition to these steric effects, *O*-GlcNAc-mediated changes in intermolecular interactions could also derive from bulk changes in the chemical properties of the LCRN. *O*-GlcNAc is uncharged, unlike phosphorylation—to which *O*-GlcNAc is often compared and which is potentially reciprocally regulated ^69^. Thus, *O*-GlcNAc weakens LCRN interactions with-out the benefit of charge-charge repulsion or attraction, which plays a critical role in how phosphorylation disrupts or enhances condensates and fibrils formed by other RNA-binding protein LCRs ^40, 41, 44^. Instead, *O*-GlcNAc moieties may disfavour phase separation by altering the hydration properties of the EWS LCR_N_ ^70^. In addition, the *O*-GlcNAc moiety may itself engage in unique intramolecular interactions with the polypeptide chain that compete with the cohesive intermolecular interactions that are important for phase separation and/or aggregation. Evidence from solid-state NMR studies of *O*-GlcNAcylated FG-Nup hydrogels suggest that such novel sugar-protein interactions do exist and are important for altering the rigidity and secondary structural propensities of FG-Nup molecules within the hydrogel ^22^.

For several RNA-binding proteins like EWS, phase-separation propensity has been found to correlate with prion-like sequence characteristics ^50, 71^. Prions are associated with their propensity to form amyloid fibrillar aggregates that have repetitive cross-beta structure and often exhibit a high resistance to disassembly via chemical or thermal denaturation ^72^. Though possessing prion-like sequence characteristics does not necessarily predispose a protein to form such stable structures under physiological conditions, there are a growing number of cases linking the phase separation of PLD-containing proteins with the emergence of fibrillar aggregate states, especially in connection with neurodegenerative disease mutations ^30, 33, 38^. It therefore seems probable that cellular mechanisms such as *O*-GlcNAcylation contribute to preventing the aggregation of PLD-containing proteins, particularly in the context of biomolecular condensates in which the nucleation of such aggregates can be promoted by high protein concentrations.

EWS was recently shown to be co-translationally *O*-GlcNAcylated, and is *O*-GlcNAcylated in several different cell lines ^54, 55, 57, 73^. These observations suggest that *O*-GlcNAcylation is a constitutive feature of EWS that imparts a persistent regulatory effect on its cellular functions. Our data provide evidence that *O*-GlcNAcylation could be important for regulating EWS aggregation in the context of cellular condensates, which include poly-ADP-ribose-induced DNA damage puncta, paraspeckles and stress granules ^64, 74, 75^.

EWS *O*-GlcNAcylation was originally discovered in the context of the chimeric fusion protein, EWS-FLI1, in which the C-terminal EWS RBD is swapped for the DNA binding domain of the Friend Leukemia Integration 1 transcription factor (FLI1) due to a chromosomal translocation ^55^. EWS-FLI1 is the signature oncogenic transcription factor in Ewing sarcoma family tumors, with the EWS LCR_N_ acting as a classical activation domain complementing the DNA-binding activity of FLI1. Global inhibition of hexosamine biosynthesis reduces EWS-FLI1 *O*-GlcNAcylation and diminishes the transcriptional output from the EWS-FLI1-targeted Id2 locus in a Ewing Sarcoma cell line ^55^. While these results are complicated by the broad off-target effects of hexosamine biosynthesis inhibition, they nevertheless provide compelling initial evidence that *O*-GlcNAcylation affects EWS-FLI1 transcriptional regulation. Recent reports have linked transcription to biomolecular condensates formed by phase separation of transcription factor and coactivator IDRs ^76, 77^. EWS-FLI1 has been hypothesized to enhance transcription by forming aberrant condensates on repetitive DNA targets of the FLI1 DNA-binding domain ^78^, supported by its colocalization to dynamic nuclear ‘hubs’ containing RNA Pol II ^49^. In light of our *in vitro* results, it would be interesting to test if *O*-GlcNAcylation of EWS-FLI1 and other elements of the transcription apparatus, such as the RNA Pol II C-terminal domain (CTD) and TATA-binding protein, affect their partitioning or activity within transcriptional condensates. Indeed, the RNA Pol II CTD and other transcription-associated factors are *O*-GlcNAcylated in the pre-initiation complex and are deglycosylated during elongation ^79, 80^. We speculate that, like phosphorylation ^81^, *O*-GlcNAcylation could regulate the exchange of RNA Pol II and other transcription-associated factors between sequential condensates associated with transcription. Further studies are needed to illuminate the rich regulatory potential of *O*-GlcNAcylation for many biomolecular condensates, including those associated with transcription, DNA repair and stress.

## EXPERIMENTAL SECTION

Unless stated otherwise, all chemicals were purchased from BioBasic Canada, Inc. (Markham, ON, Canada).

### Plasmids and Cloning

The coding sequence for *Homo sapiens EWSR1*, residues 1-264 (*N*-terminal LCR; LCR_N_) was synthesized by GenScript (Piscataway, NJ, USA) with codon optimization for expression in *Escherichia coli*. The LCR_N_ coding sequence was inserted in frame with a *N*-terminal hexahistidine (His_6_)-SUMO tag in a Champion pET SUMO expression vector (Invitrogen, Carlsbad, CA, USA) by Gibson Assembly (New England Biolabs, Ipswich, MA, USA). The coding sequence for EWS 265-656 (the RNA binding domains; RBD) was derived from *H. sapiens EWSR1* cDNA obtained from the NIH Mammalian Gene Collection (MGC), made available through the SickKids SPARC cDNA archive (https://lab.research.sickkids.ca/sparc-drug-discovery/services/molecular-archives/sparc-cdna-archive/). The RBD fragment was amplified via polymerase chain reaction using Kapa HiFi HotStart ReadyMix (Roche, Basel, Switzerland) and then Gibson assembled in frame with an *N*-terminal His_6_-SUMO tag in the same vector as above. This construct contains a diglycine insertion at the *N*-terminus of the EWS segment (i.e. directly following the SUMO tag and before residue 265 of EWS).

The pET24b *H. sapiens* ^(nucleocytoplasmic variant, 110kDa) *E. coli* expression plasmid was a kind gift from Suzanne Walker’s lab. The K842M mutation was introduced via Site-Directed Mutagenesis using a QuikChange II Kit (Agilent, Santa Clara, CA, USA).

### Recombinant Protein Expression

The pET SUMO EWS LCR_N_ and EWS RBD plasmids were transformed into *E. coli* BL21-CodonPlus (DE3) RIPL chemically competent cells (Agilent). EWS RBD methylation was achieved by cotransformation along with a glutathione *S*-transferase-linked protein arginine methyltransferase 1 (*PRMT1*) *E. coli* expression vector, as previously described ^82^. Successful transformants were selected on lysogeny broth (LB)-agar plates containing 50µg mL^-1^ kanamycin sulfate and 34µg mL^-1^ chloramphenicol (also including 100µg mL^-1^ ampicillin in the case of PRMT1 co-expression). Single colonies were used to inoculate ∼50mL LB overnight cultures containing the same antibiotics. Following overnight incubation at 37°C with 250rpm shaking, these starter cultures were used to inoculate liter-scale cultures which were then incubated under the same conditions until an optical density at (OD_600_) of ∼1.0 was attained. Recombinant protein expression was initiated by the addition of 0.5mM isopropyl-β-D-thiogalactopyranoside (IPTG). The cultures were then incubated for another ∼18 hours at 20°C prior to being harvested by centrifugation in a JLA9.1000 rotor operating in an Avanti J-26 XP centrifuge (Beckman Coulter, Brea, CA, USA). Cell pellets were suspended in lysis buffer (25mM Tris pH 8.0, 4M guanidinium chloride, 500mM NaCl, 20mM imidazole (Millipore-Sigma Canada, Oakville, ON, Canada), 5mM 2-mercaptoethanol (Millipore-Sigma), pH 8.0) and stored at -20°C until further use.

### Protein Purification

Cells were lysed by sonication using a QSonica Q500 sonicator outfitted with a 6mm probe operating at 25% amplitude with 2s pulses / 50% duty for 8 min. total. Lysates were clarified by centrifugation in a JA-20 fixed-angle rotor (Beckman Coulter) operated at 15000 rpm for 1 hour at 4°C. The supernatant was loaded onto a gravity column containing a ∼10mL bed volume of Ni Sepharose resin (Cytiva, Marlborough, MA, USA) pre-equilibrated in lysis buffer. After repeated washes with lysis buffer, the bound proteins were eluted with lysis buffer containing 280mM imidazole. These fractions were pooled and dialyzed against ULP buffer (20mM HEPES, 150mM NaCl, 2mM 2-mercaptoethanol) using a 3kDa molecular weight cut-off (MWCO) regenerated cellulose membrane (Spectrum Chemical MFG Co., New Brunswick, NJ, USA). His_6_-Ulp1 protease (purified in-house) was then added to cleave the SUMO-tag during overnight dialysis at 4°C. The protease-treated samples were re-loaded over the nickel column to separate the His_6_-SUMO tag and protease from the cleaved protein. The flow-through was concentrated using an Amicon 15mL-volume centrifugal concentrator unit equipped with a 3kDa MWCO membrane (EMD-Millipore, Burlington, MA, USA) before being injected onto a Superdex 75 16/600 HiLoad column (Cytiva) pre-equilibrated in size exclusion chromatography (SEC) buffer (25mM Tris, 2M guanidinium chloride, 2mM 2-mercaptoethanol, pH 7.5) on an AKTA FPLC chromatography system (Amersham Biosciences Co., Little Chalfont, UK). Proteins were eluted isocratically at a constant flow rate of 0.5mL min^-1^ with fractions collected every 2 min. Fractions were analyzed by SDS-PAGE with Coomassie staining.

At this stage, the EWS LCR_N_ was sufficiently pure for further experiments, whereas the RBD required an additional ion exchange purification step to remove excess impurities. The pooled RBD fractions (with or without PRMT1-methylation) were pooled and dialyzed against 25mM HEPES, 50mM NaCl, 0.5mM EDTA, 2mM 2-mercaptoethanol, pH 6.5 in a 3kDa MWCO membrane. The dialyzed sample was clarified by centrifugation at 7000rpm in a fixed-angle Thermo F13 rotor in a Sorvall Legend XFR centrifuge (Thermo Scientific, Waltham, MA, USA) at 4degC for 25 minutes before being loaded onto a 5mL HiTrap sulfopropyl (SP) HP column (Cytiva) pre-equilibrated in the same dialysis buffer as above. After washing the resin with several column volumes of dialysis buffer, the bound protein was eluted over a buffer gradient with increasing NaCl concentration ranging from 50mM to 2M (buffer components were otherwise the same as in the dialysis buffer). Fractions were analyzed by SDS-PAGE; most of the RBD eluted at salt concentrations exceeding 0.5M. The purest fractions were pooled and flash frozen for storage at -80°C.

Wild-type and K842M OGT were purified as previously described ^83^ with slight modifications. Following the nickel purification, OGT samples were dialyzed against 25mM Tris, 50mM NaCl, 0.5mM EDTA, 2mM dithiothreitol (DTT), pH 7.5 and then concentrated using a centrifugal concentrator equipped with a 50kDa MWCO membrane. Concentrated samples were injected onto a Superdex 200 16/600 HiLoad column (Cytiva) pre-equilibrated in the same dialysis buffer on an AKTA FPLC chromatography system (Amersham Biosciences Co.). The system was run at 4°C with a constant flow rate of 0.5mL min^-1^ with fractions collected every 2min. Fractions were analyzed by SDS-PAGE and Coomassie staining to assess purity. The protein was concentrated to working concentrations of ∼20-25µM before being flash frozen and stored at -80°C until further use.

### *In vitro O*-GlcNAcylation

Unless specified otherwise, *O*-GlcNAcylation reactions contained 10µM of the appropriate protein substrate (e.g., EWS LCR_N_, RBD), 1µM OGT (wild-type or K842M), 1mM UDP-GlcNAc (Millipore-Sigma), 25mM Tris, 50mM NaCl, 0.5mM EDTA, 2mM dithiothreitol, pH 7.5. The sample volume was scaled according to the quantity of *O*-GlcNAcylated product desired. Analytical-and preparative-scale reactions were incubated for 14 hours at laboratory temperature (∼22°C) with gentle rocking. Analytical-scale samples were prepared directly for mass spectrometry at this stage (see below), while preparative-scale samples were quenched by the addition of guanidinium chloride (to 2M final) to facilitate purification by SEC. Concentrated samples were injected onto a Superdex 75 16/600 Hi-Load column pre-equilibrated in 25mM Tris, 2M guanidinium chloride, 2mM 2-mercaptoethanol, pH 7.5 on an AKTA FPLC chromatography system. Fractions containing *O*-GlcNAcylated protein were analyzed by SDS-PAGE (a slight migration shift due to *O*-GlcNAcylation was visible as compared to the unmodified protein) and mass spectrometry. The protein was concentrated to ∼150µM before being flash frozen and stored at -80°C.

### Intact Mass Spectrometry (MS)

All MS experiments were conducted at the Structural Genomics Consortium (SGC) Toronto facility. Samples were either prepared directly from analytical-scale *O*-GlcNAc reactions or from preparative-scale purified samples that were first exchanged out of guanidinium chloride-containing buffer into 25mM Tris, pH 7.5. In either case, formic acid was added to a final concentration of 0.1%v/v. 2µL of each sample were injected onto an Agilent UPLC-quadrupolar time of flight (Q-ToF) 6545 MS system equipped with a Dual AJS electrospray ionization source operating in positive ion mode. Samples were desalted online through a C18 column into a mobile phase containing 95%v/v acetonitrile, 4.5%v/v H_2_O, 0.5%v/v formic acid. Spectra were recorded over a scan range of 500-3200 m/z at a time interval of 1 scan per second. Raw spectra were processed in Agilent Masshunter software and deconvoluted using the maximum entropy algorithm over a mass range of 10-50kDa with 1Da resolution. Raw and deconvoluted spectra were plotted in GraphPad Prism 6.

### Fluorescent Protein Labelling

Aliquots of unmodified or *O*-GlcNAcylated EWS LCR_N_ were dialyzed in 3.5kDa MWCO GebaFlex midi cassettes (Gene Bio-Applications Ltd., Yavne, Israel) against 25mM sodium phosphate, 2M guanidinium chloride, pH 6.5, to remove the Tris prior to labelling. Sulfo-cyanine-3 dye conjugated to *N*-hydroxysuccinimide (sulfo-Cy3 NHS-ester; Lumiprobe Co., Hunt Valley, MD, USA) was dissolved in dimethylformamide (Sigma) to a working concentration of 20mM. An appropriate volume of the dye stock was added into a 20µM sample of dialyzed protein to yield a final dye concentration of 200µM. This mixture was allowed to incubate overnight in the dark at 4°C to promote selective labelling of the *N*-terminus. The reaction was quenched by the addition of 25mM Tris, pH 7.5 and filtered through a 0.22µm polyether-sulfone membrane (GE Healthcare, Chicago, IL, USA) to remove any aggregates. The filtrate was desalted into 25mM Tris, 2M guanidinium chloride, pH 7.5 using a 5mL HiTrap desalting column (Cytiva) running at a constant flow rate of 1.5mL min^-1^ on an AKTA FPLC system. Fractions containing the labelled protein were analyzed by SDS-PAGE to ensure that fluorescent reaction by-products had been removed. Intact mass spectrometric analysis was also used to confirm labelling. Polyacrylamide gels containing the sulfo-Cy3-modified protein were visualized on a BioRad Chemidoc MP system with Cy3 emission/excitation settings.

### Droplet Formation and Microscopy

For all phase separation experiments, EWS LCR_N_ (with or without *O*-GlcNAcylation or sulfo-Cy3-labelling) was exchanged into 25mM Tris, pH 7.5, by successively concentrating and diluting the protein out of the original guanidinium chloride-containing storage buffer. During this process, the temperature of the centrifuge chamber was set to ≥37°C to limit precipitation which occurred more readily at lower temperature. Droplet formation was induced by combining protein samples 1:1 with buffers containing twice the appropriate NaCl concentration at room temperature. For DIC microscopy, droplet samples were pipetted directly into the wells of 96-well polystyrene glass-bottom plates (Eppendorf, Hamburg, Germany). Images were acquired on a Zeiss Axio Observer 7 microscope with a 40x air objective lens. For fluorescence microscopy, 5µL samples were pipetted into 35mm-diameter uncoated no. 1.5 glass-bottom dishes (MatTek, Ashland, MA, USA) which were subsequently sealed with a glass coverslip to limit evaporation. To enable non-wetting conditions for fusion experiments, the dishes were pre-treated with Sigmacote (Millipore-Sigma) according to the manufacturer’s instructions. Fluorescence imaging was performed on a Leica TCS SP8 Lightning / DMi8 system equipped with a Hamamatsu C9100-13 EM-CCD camera. All images were acquired with a 63x oil-immersion objective lens at 1024×1024 pixel resolution. Sulfo-Cy3 fluorescence was detected with a Leica hybrid detector (HyD) following excitation with a 561nm (40mW) laser. Image files were analyzed and processed using Fiji (https://imagej.net/Fiji).

### Fluorescence Recovery After Photobleaching (FRAP)

FRAP data were collected using the LAS X FRAP module on the same Leica system described above. Circular, 2.5µm-diameter regions of interest (ROI) were positioned in the centres of droplets for bleaching, which was performed in a single pulse at 30% laser power on zoom-in mode. After photobleaching, images were acquired at a frequency of 0.77s^-1^. Fluorescence intensity measurements from inside the bleached ROI were normalized to measurements from a separate ROI in an adjacent unbleached droplet to correct for passive photobleaching and focus drift. The values were then normalized to the average of 5 pre-bleach scans to yield relative intensity values. The recovery timescale was obtained by fitting a single exponential curve to the processed data: *I*(*t*) =*a*− *be*^−*t*/*τ*^, where *I* (t) is relative intensity at each time, t, and a, b and *τ*are fit parameters. Half-times were calculated as *t*_1/2_ = *τ*In 2.

### Turbidity Assays

Samples were prepared in 96-well clear polystyrene plates prior to being transferred into a 384-well black polystyrene glass-bottom plate (Corning, Corning, NY, USA) for measurement on a SpectriMax i3x plate reader (Molecular Devices, San Jose, CA, USA). The assay buffers typically contained 25mM Tris, 0.5mM EDTA, 2mM DTT, pH 7.5 with a variable NaCl concentrations (to promote or prevent droplet formation) depending on the experimental setup. All sample had a final volume of 15 µL per well. Absorbance measurements were collected at 25 with 500nm-wavelength light at a read-height of 12mm. Data were processed and plotted in GraphPad Prism 6.

### Temperature Ramp Experiments

EWS LCR_N_ (with or without *O*-GlcNAcylation) was dialyzed against 20mM 3-(*N*-morpholino)propanesulfonic acid (MOPS), pH 7.5 in a 3kDa MWCO GebaFlex midi cassettes (Gene Bio-Applications Ltd.) overnight at room temperature. 200µL samples were prepared at 30µM final protein concentration in 20mM MOPS, 150mM NaCl, pH 7.5, with different ratios of unmodified:modified EWS LCR_N_ present. Samples were pipetted into 250µL 1mm-thick quartz cuvette for turbidity measurements in a JASCO J-1500 circular dichroism spectrophotometer.

### Calculating percentages of predicted phase separating proteins

The predictions of phase separation propensities for human proteins were available from Vernon *et al*. ^67^. The predictions from three different metrics (catGRANULE ^65^, PScore ^9^ and PLAAC ^66^) were considered, with method-specific cut-offs used to define propensity to phase separate in a binary manner. A value larger than zero for catGRAN-ULE score was used to define propensity to phase separate. The same cut-off (>0) was used to identify proteins that contain probable prion-like domains by PLAAC. The cut-off of larger or equal than four was used for PScore, as previously established by the authors of the original work. The prediction scores of the three metrics were in addition expressed as a percentage of the human proteome predicted to phase separate in order to unify different scoring schemes across the predictors, with 4% of the human proteome used as a cut-off. The list of sequences of all proteins currently in the PhosphositePlus database was retrieved from the website (https://www.phosphosite.org/hosphosite_seq.fasta). The set of *O*-GlcNAc modified protein sequences was retrieved from the same database (O-GlcNAc_site_dataset). Both datasets were filtered to include human proteins only. The fractions of the proteins predicted to phase separate in each of the categories: i) proteome; ii) PhosphositePlus; and iii) *O*-GlcNAc, were computed by dividing the number of proteins above the cut-off of each of the metrics (PScore, PLAAC, catGRANULE) by the total number of proteins in each category. A Fisher’s exact test was used to assess the significance of the difference between the fraction of proteins above the threshold in the human proteome and each of the categories (the *p*-values of the test given in Table S1; and significance indicated with i) ‘*’ *P* ≤0.01; ii) ‘**’ *P* ≤0.001; iii) ‘***’ *P* ≤0.0001).

### Defining proteins with long IDRs

SPOT-Disorder ^84^ was run on the reference human proteome from Uniprot (UP000005640, downloaded on Aug 8, 2019), to obtain a residue-level prediction of intrinsic disorder for all human proteins (canonical sequences only). Short sequences of seven or less residues of predicted ‘order’ mapped in between long predicted disorder regions were concatenated into the predicted disordered regions. A cut-off of 100 consecutive residues of predicted disorder was used to define ‘long IDRs’.

### Cell culture and siRNA transfection

Please refer to Table S2 for a detailed list of antibodies used in this study. HeLa cells were cultured in Dulbecco’s Modified Eagle Medium (DMEM) with 10% fetal bovine serum and 1% penicillin-streptomycin. To downregulate expression of *OGT*, cells were reverse-transfected with 20 nM ON-TARGETplus Human *OGT* siRNA (Horizon, cat. no. J-019111-06) for 48 hours using INTERFERin siRNA Transfection Reagent (Polyplus, Illkirch-Graffenstaden, France). Knockdown efficiency was verified by immunoblotting with an anti-OGT antibody (Abcam, Cambridge, UK), while global *O*-GlcNAc levels were assessed by immunoblotting with an anti-*O*-GlcNAc (RL2) antibody (Abcam).

*O*-GlcNAc stoichiometries on EWS and TAF15 in HeLa cell lysates were measured by Western blotting with the appropriate antibodies (Abcam) after the lysates were processed with the ClickIT *O*-GlcNAc Enzymatic Labelling and ClickIT detection assay kits (Invitrogen). Volumes of lysate containing ∼200µg of protein dissolved in FRA buffer (see below) were used for these assays. The manufacturer’s instructions were followed, with the exception that the alkyne-biotin detection reagent from the latter kit was replaced with methoxy-PEG-alkyne (average MW ∼5kDa; Biochempeg, Watertown, MA, USA), which was present at a final concentration of 5mM during the Click chemistry step. After the final protein precipitation step, the pellets were resuspended in 50µL of 4x Bolt LDS sample buffer (Invitrogen) after vigorous vortexing and heating at 70°C. Samples were diluted four-fold with water before being loaded onto a 4-20% tris-glycine polyacrylamide gel (BioRad, Hercules, CA, USA), approximately ∼10µg of protein (i.e., 4µL of undiluted dissolved pellet diluted to 16µL with water) per lane. SDS-PAGE and Western blotting were performed according to standard protocols.

### Filter retardation assay (FRA)

siRNA-transfected cells were harvested and lysed in FRA lysis buffer (20 mM Tris-HCl pH 7.5, 150 mM NaCl, 1 mM EDTA, 1 mM EGTA, 1% Triton X-100, 1x protease inhibitor cocktail) and protein concentration was quantified using a Pierce 660 assay (Thermo Scientific, cat. no. 22660). 10µg of lysate was loaded onto a methanol-activated 0.2µm cellulose-acetate membrane in a BioRad slot-blot apparatus. The wells of the apparatus were washed prior to and after sample loading using RIPA buffer (50mM Tris-HCl pH 7.5, 150mM NaCl, 1%v/v Triton X-100, 0.5%w/v sodium deoxycholate, 0.1%w/v SDS, 5 mM EDTA, 1x protease inhibitor cocktail). Following sample loading and washes, the cellulose-acetate membrane was fixed in methanol for 5 minutes, washed in trisbuffered saline with 0.1%v/v Tween-20 (TBST) for 5 minutes, and subsequently processed for immunoblotting. In brief, the membrane was blocked in 5%w/v skim milk powder / TBST, incubated in primary antibody (anti-EWS and anti-TAF15; overnight at 4°C), incubated in horseradish peroxidase (HRP)-linked secondary antibody (anti-rabbit IgG; 1 hr at RT), and developed on a ChemiDoc Gel Imaging System using Luminata Crescendo Western HRP Substrate (Sigma, WBLUR0500). Signal intensity was quantified using the Gel Analyzer plugin in ImageJ. Three technical replicates were performed for each experiment. Western blots were performed in parallel to ensure equal loading and levels of expression of the proteins of interest.

## ASSOCIATED CONTENT

### Supporting Information

Supplementary mass spectrometry data, including raw spectra of unmodified and *O*-GlcNAcylated EWS LCR_N_ (Figure S1). Fluorescence micrographs to accompany FRAP data from LCR_N_+RBD droplets in Fig. 4 (Figure S2). Western blots showing (i) depletion of OGT following siOGT treatment of HeLa cells, (ii) lack of *O*-GlcNAcylation on TAF15 in HeLa cells, (iii) negligible changes in EWS and TAF15 expression levels following siOGT treatment (Figure S3). Supplementary bioinformatics data on differential prediction of phase separation propensity for proteins in the human proteome, including those filtered for containing long (>100 residue) IDRs (Figure S4). Tabulated data corresponding to the bioinformatic analyses presented in Figs. 6 and S4 (Table S1). A listed of antibodies used in this study (Table S2).

The Supporting Information is available free of charge on the ACS Publications website.

## AUTHOR INFORMATION

### Notes

The authors declare no competing financial interest.

### Funding Sources

M.N. is funded by an Alexander Graham Bell Canada Graduate Scholarship—Doctoral from the Natural Sciences and Engineering Research Council (NSERC) and M.T. by a Vanier Graduate Scholarship. This work was supported by funding to J.D.F.-K. from the Canadian Institutes for Health Research (CIHR) FDN-148375 and the Canadian Cancer Society, to both J.D.F.-K. and H.O.L. from the Canada Research Chair program, and to H.O.L. from NSERC and University of Toronto’s Connaught Fund and Medicine by Design, which receives funding from the Canada First Research Excellence Fund. Additional support was from Children’s Cancer Foundation, Inc. as a Force 3 Fellowship (J. A.T.), and the NIH R01CA233619-01A1 (J.A.T.), LCCC CCSG Grant P30 CA051008-16 (Lou Weiner, PI, J.A.T.).

## ACKNOWLEDGMENT

We would like to thank the SickKids Imaging Facility for their assistance with the microscopy experiments.

## Notes

### Competing Interest Statement

The authors have declared no competing interest.

