## Supplemental Information for "*O*-GlcNAcylation reduces phase separation and aggregation of the EWS N-terminal low complexity region"

Supplementary Figures S1-S4, Supplementary Tables S1 & S2.

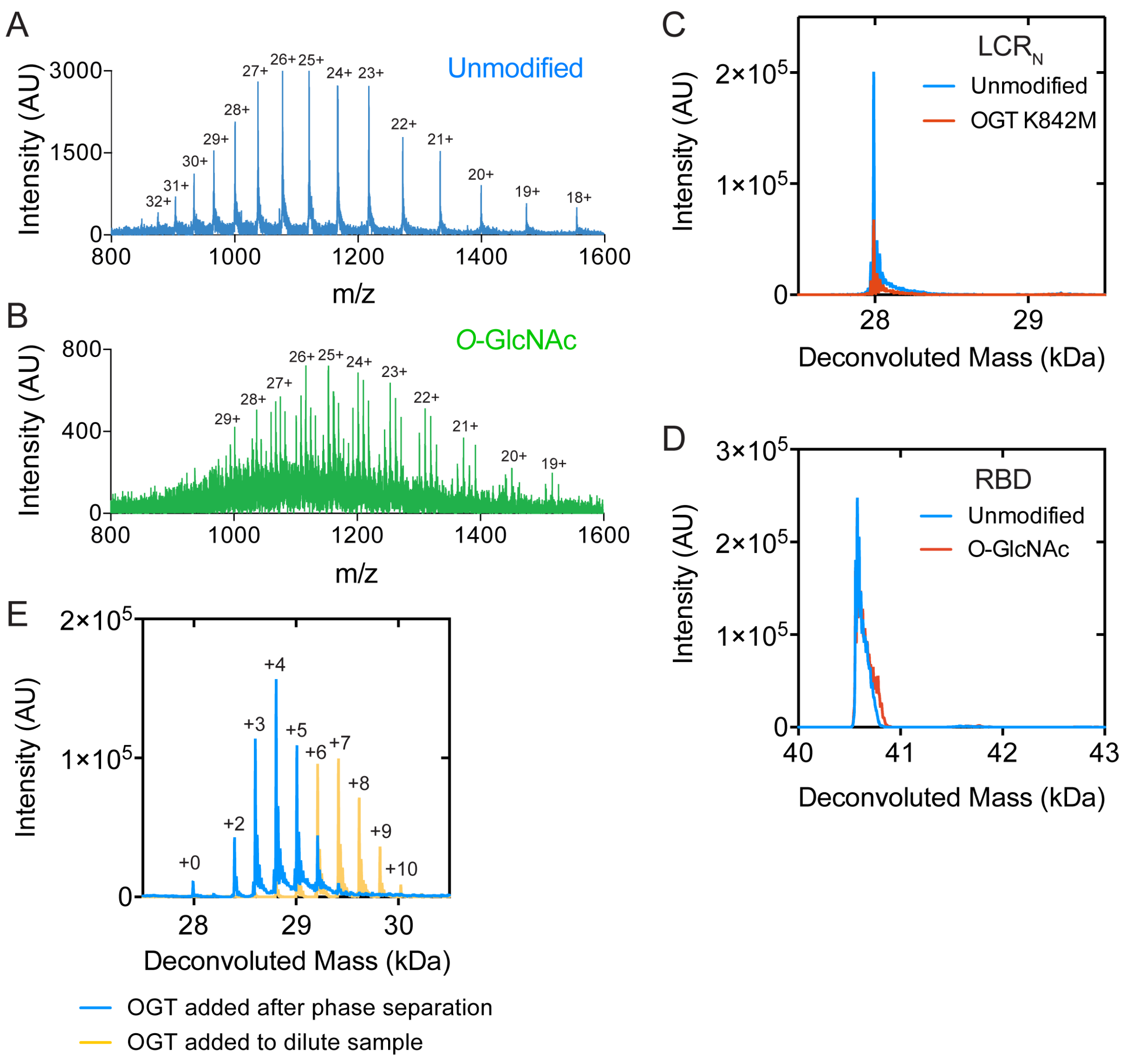


**Figure S1.** Mass spectrometry data for EWS $LCR_N$. (A) Mass spectra of unmodified and (B) *O*-GlcNAcylated EWS $LCR_N$. Numbers indicate charge state assignment. (C) Deconvoluted mass spectrum of EWS $LCR_N$ before and after modification by OGT K842M. (D) Deconvoluted mass spectrum of EWS RBD before and after modification by OGT. Calculated molecular weight: 40,618.07 Da. (E) Deconvoluted mass spectra of EWS $LCR_N$ modified by OGT following phase-separation at 200mM NaCl and in a dilute sample without inducing phase separation.

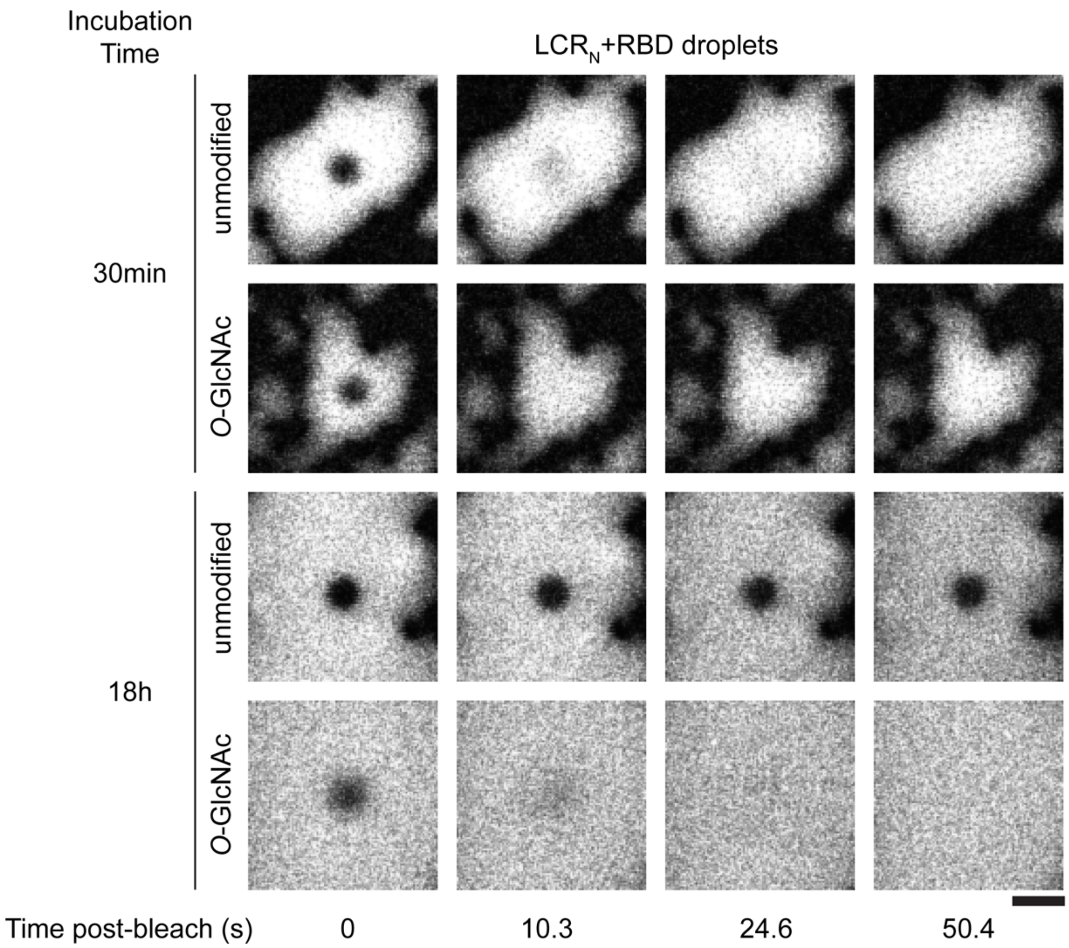


**Figure S2.** Representative fluorescence micrographs of EWS LCR$_N$+RBD FRAP time points at early and late incubation periods. All samples contain 10μM LCR$_N$ (with or without *O*-GlcNAc), 100nM sulfo-Cy3-labelled LCR$_N$ (with or without *O*-GlcNAc), 10μM RBD, 25mM Tris, 100mM NaCl, 0.5mM tetrasodium EDTA, 2mM dithiothreitol, pH 7.5 at lab temperature. Scale: 5μm.

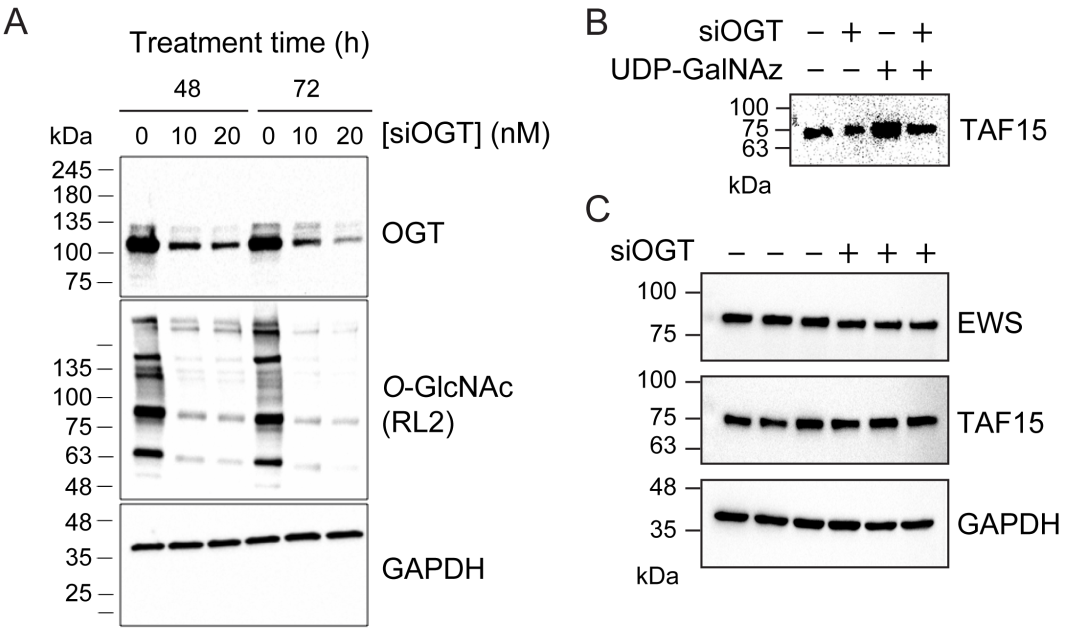


**Figure S3.** Changes in protein expression and *O*-GlcNAylation following siOGT treatment in HeLa cells. (A) Western blots showing diminished *OGT* expression and general *O*-GlcNAcylation levels in HeLa cells following siOGT knockdown compared to GAPDH. Treatment time indicates the duration with which the cells were exposed to growth media containing siOGT. The 48-hour incubation, 20nM condition was selected for the remaining experiments (e.g., the FRA). (B) Western blot of chemoenzymatic (ClickIT) labelling of potential *O*-GlcNAc sites demonstrating that TAF15 is unmodified. UDP-GalNAz indicates whether O-GlcNAc moieties were primed with GalNAz for covalent addition of mPEG-alkyne. (C) Representative Western blots of FRA sample lysates, showing similar EWS, TAF15 and GAPDH expression levels regardless of siOGT treatment. GAPDH, glyceraldehyde 3-phosphate dehydrogenase.

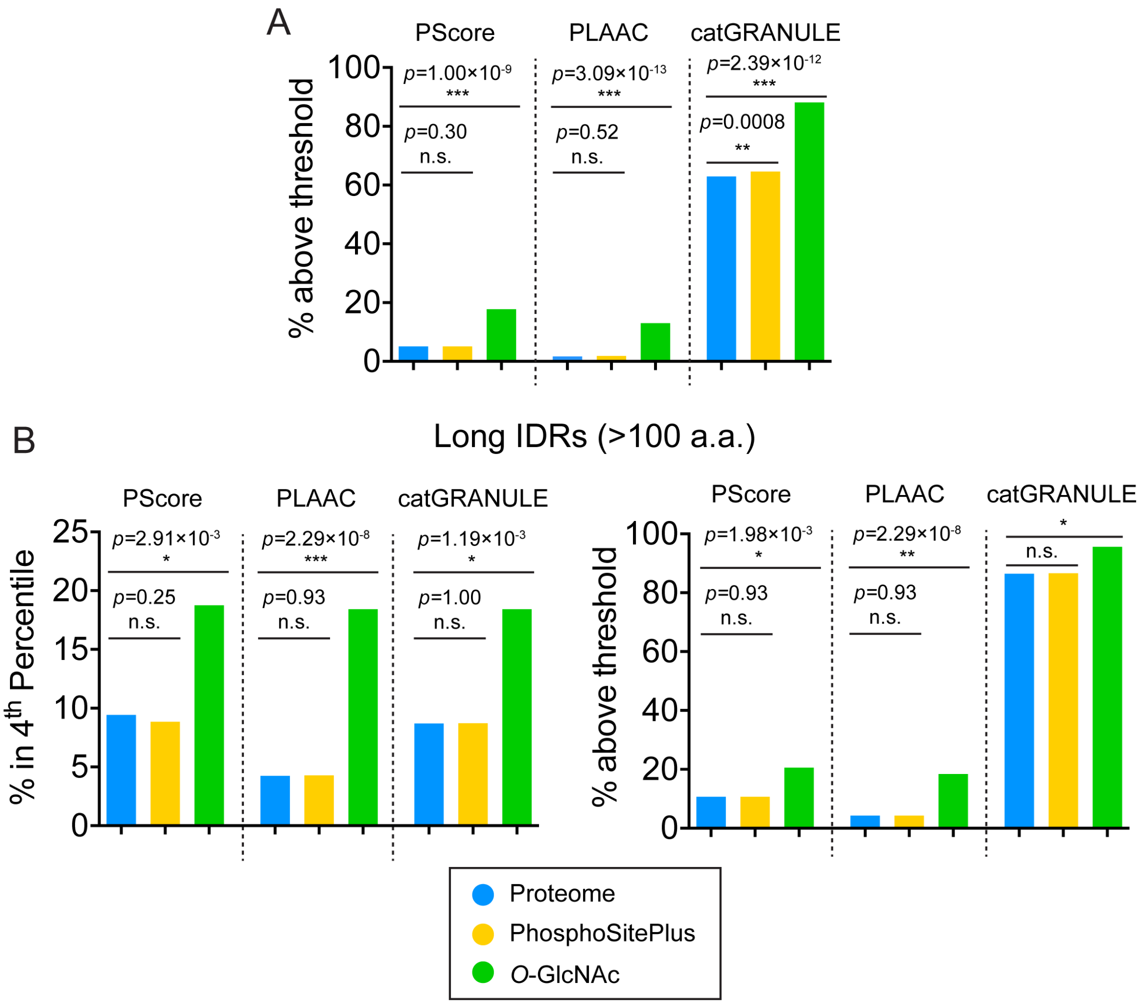


**Figure S4.** Supplementary bioinformatics data on differential prediction of phase separation for proteins in the human proteome, in the PhosphoSitePlus database and PhosphoSitePlus-annotated *O*-GlcNAcylated proteins. (A) Comparison of proteins scoring above the method-specific threshold by three different metrics of phase separation propensity (PScore, PLAAC and catGRANULE, see Methods). (B) Comparison of proteins with at least one predicted intrinsically disordered region spanning one hundred residues or more (defined as 'long IDRs', see Methods) according to their scoring within the top 4$^{th}$ percentile (left) or above the method-specific threshold (right) of the three phase-separation predictive metrics. Indicated *p*-values were derived from Fisher's exact test.

**Supplementary Tables**

**Table S1.** Comparative analysis of predicted phase-separating propensities using PScore, PLAAC and catGRANULE across the human proteome, PhosphoSitePlus database, and PhosphoSitePlus-annotated *O*-GlcNAcylated proteins. The top panel provides values for the metric-specific score thresholds for phase separation (PScore: ≥4, PLAAC: >0, catGRANULE: >0). The bottom panel provides values for the top 4th percentile for each metric.

| **PScore** | | **Total** | **With Score** | **Score ≥4** | ***P*-val with respect to Proteome** | **With Predictor Score and IDR>100aa** | **With Predictor Score ≥4 and IDR>100aa** | ***P*-val with respect to Proteome** |
|---|---|---|---|---|---|---|---|---|
| | Proteome | 21047 | 18455 | 943 | N/A | 6906 | 736 | N/A |
| | PhosphoSitePlus | 19990 | 18286 | 939 | 0.2989 | 6869 | 736 | 0.9340 |
| | *O*-GlcNAc | 171 | 163 | 29 | 1.00E-09 | 112 | 23 | 1.98E-03 |

| **PLAAC** | | **Total** | **With Score** | **Score >0** | ***P*-val with respect to Proteome** | **With Predictor Score and IDR>100aa** | **With Predictor Score >0 and IDR>100aa** | ***P*-val with respect to Proteome** |
|---|---|---|---|---|---|---|---|---|
| | Proteome | 21047 | 21047 | 353 | N/A | 7046 | 299 | N/A |
| | PhosphoSitePlus | 19990 | 19990 | 352 | 0.5200 | 6977 | 299 | 0.9300 |
| | *O*-GlcNAc | 171 | 169 | 22 | 3.09E-13 | 114 | 21 | 2.29E-08 |

| **catGRANULE** | | **Total** | **With Score** | **Score >0** | ***P*-val with respect to Proteome** | **With Predictor Score and IDR>100aa** | **With Predictor Score >0 and IDR>100aa** | ***P*-val with respect to Proteome** |
|---|---|---|---|---|---|---|---|---|
| | Proteome | 21047 | 21047 | 13241 | N/A | 7046 | 6092 | N/A |
| | PhosphoSitePlus | 19990 | 19990 | 12895 | 0.000785 | 6977 | 6046 | 0.7476 |
| | *O*-GlcNAc | 171 | 169 | 149 | 2.39E-12 | 114 | 109 | 2.12E-03 |

| **PScore** | | **Total** | **With Score** | **With Score in top 4th percentile** | ***P*-val with respect to Proteome** | **With Predictor Score and IDR>100aa** | **With Score in top 4th percentile and IDR>100aa** | ***P*-val with respect to Proteome** |
|---|---|---|---|---|---|---|---|---|
| | Proteome | 21047 | 18455 | 841 | N/A | 6906 | 651 | N/A |
| | PhosphoSitePlus | 19990 | 18286 | 823 | 0.5479 | 6869 | 608 | 0.2489 |
| | *O*-GlcNAc | 171 | 163 | 26 | 7.39E-09 | 112 | 21 | 2.91E-03 |

| **PLAAC** | | **Total** | **With Score** | **With Score in top 4th percentile** | ***P*-val with respect to Proteome** | **With Predictor Score and IDR>100aa** | **With Score in top 4th percentile and IDR>100aa** | ***P*-val with respect to Proteome** |
|---|---|---|---|---|---|---|---|---|
| | Proteome | 21047 | 21047 | 353 | N/A | 7046 | 299 | N/A |
| | PhosphoSitePlus | 19990 | 19990 | 352 | 0.5185 | 6977 | 299 | 0.9334 |
| | *O*-GlcNAc | 171 | 169 | 22 | 3.94E-13 | 114 | 21 | 2.29E-08 |

| catGRANULE | | Total | With Score | With Score in top 4th percentile | *P*-val with respect to Proteome | With Predictor Score and IDR>100aa | With Score in top 4th percentile and IDR>100aa | *P*-val with respect to Proteome |
|---|---|---|---|---|---|---|---|---|
| | Proteome | 21047 | 21047 | 841 | N/A | 7046 | 613 | N/A |
| | PhosphoSitePlus | 19990 | 19990 | 823 | 0.5480 | 6977 | 608 | 1.000 |
| | *O*-GlcNAc | 171 | 169 | 26 | 7.39E-09 | 114 | 21 | 1.19E-03 |

**Table S2.** Antibodies used in this study**.**

| **Antigen (Notes)** | **Species, isotype** | **Supplier** | **Serial Number** | **Dilution Used (Western Blotting, FRA)** |
|---|---|---|---|---|
| **EWS** | Rabbit, IgG | Abcam | ab133288 | 1:10000 |
| **TAF15** | Rabbit, IgG | Abcam | ab134916 | 1:10000 |
| **OGT** | Rabbit, IgG | Abcam | ab96718 | 1:1000 |
| ***O*-GlcNAc (RL2)** | Mouse, IgG | Abcam | ab2739 | 1:5000 |
| **GAPDH** | Rabbit, IgG | Cell Signaling | 5174S | 1:5000 |
| **Rabbit IgG (horseradish peroxidase-linked)** | Goat | Cell Signaling | 7074V | 1:10000 |
| **Mouse IgG (horseradish peroxidase-linked)** | Horse | Cell Signaling | 7076V | 1:10000 |
